# Replication in the presence of dengue convalescent serum impacts Zika virus neutralization sensitivity and fitness

**DOI:** 10.1101/2022.12.21.521484

**Authors:** Jeffrey M. Marano, James Weger-Lucarelli

**Author notes:** **Correspondence:** James Weger-Lucarelli.

## Abstract

Flaviviruses like dengue virus (DENV) and Zika virus (ZIKV) are mosquito-borne viruses that cause febrile, hemorrhagic, and neurological diseases in humans, resulting in 400 million infections annually. Due to their co-circulation in many parts of the world, flaviviruses must replicate in the presence of pre-existing adaptive immune responses targeted at serologically closely related pathogens, which can provide protection or enhance disease. However, the impact of pre-existing cross-reactive immunity as a driver of flavivirus evolution, and subsequently the implications on the emergence of immune escape variants, is poorly understood. Therefore, we investigated how replication in the presence of convalescent dengue serum drives ZIKV evolution. We used an *in vitro* directed evolution system, passaging ZIKV in the presence of serum from humans previously infected with DENV (anti-DENV) or serum from DENV-naïve patients (control serum). Following five passages in the presence of serum, we performed next-generation sequencing to identify mutations that arose during passaging. We studied two non-synonymous mutations found in the anti-DENV passaged population (E-V355I and NS1-T139A) by generating individual ZIKV mutants and assessing fitness in mammalian cells and live mosquitoes, as well as their sensitivity to antibody neutralization. Both viruses had increased fitness in Vero cells with and without the addition of anti-DENV serum and in human lung epithelial and monocyte cells. In Aedes aegypti mosquitoes—using blood meals with and without anti-DENV serum—the mutant viruses had significantly reduced fitness compared to wild-type ZIKV. These results align with the trade-off hypothesis of constrained mosquito-borne virus evolution. Notably, only the NS1-T139A mutation escaped neutralization, while E-V335I demonstrated enhanced neutralization sensitivity to neutralization by anti-DENV serum, indicating that neutralization escape is not necessary for viruses passaged under cross-reactive immune pressures. Future studies are needed to assess cross-reactive immune selection in humans and relevant animal models or with different flaviviruses.

## Introduction

Dengue virus (DENV; Genus *Flavivirus;* Family *Flaviviridae)* infects roughly five percent of the global population annually [1]. Zika virus (ZIKV; Genus *Flavivirus;* Family *Flaviviridae)* emerged in the western hemisphere in 2013 [2] and is estimated to have caused over 100 million infections by 2018 [3]. ZIKV causes severe pathologies, including microcephaly and seizures (reviewed in [4]). One proposed driver of severe disease is pre-existing immunity against DENV [5–11], four genetically and serologically closely related viruses [12]. While pre-existing cross-reactive DENV immunity from antibodies can be protective [5, 8, 13], it can also enhance disease [5–11] upon ZIKV infection. Given that these cross-reactive humoral responses play a significant role in disease, their role in ZIKV evolution should also be examined to more fully understand flavivirus evolution.

Immune-driven evolution occurs when the host immune response neutralizes only a subset of viruses, placing selective pressure on the virus population [14]; the surviving viruses—which likely have some resistance to the immune pressure—become founders for a subsequent generation [15, 16]. Evolution driven by antibodies has been described for several viruses, including West Nile virus [17], Nipah virus [18], chikungunya virus [19], influenza [20–24], SARS-CoV-2 [25–27], and many others [28–33]. While attempts have been made to study the impact of cross-reactive immune-driven evolution in ZIKV, these previous studies use monoclonal antibodies [34], which are a simplistic model for the complex polyclonal human antibody response or study mutations that were not specific to immune selection[35]. It is critical that cross-reactive immune-driven evolution be studied since mutations that arise may have implications for transmission [36, 37] or disease severity [36, 38–40].

To address this gap, we examined the effects of cross-reactive antibody selection by passaging ZIKV in the presence of serum from convalescent dengue patients from the Dominican Republic or control serum from dengue-naïve donors from the United States. After passaging, we sequenced the viral populations using next-generation sequencing (NGS). Compared to the virus passaged in the control serum, the premembrane (prM) region of the anti-DENV serum passaged virus was less divergent from the starting virus and had lower non-synonymous diversity. We then examined the anti-DENV serum passaged virus for enriched mutations and engineered two unique mutations using a reverse genetics system. We assessed the impact of these two mutations, E-V355I and NS1-T139A, on fitness in various mammalian cell lines and *Aedes aegypti* mosquitoes and their sensitivity to neutralization by anti-DENV immune serum. Notably, both mutations had increased fitness in mammalian cell culture and reduced fitness in live mosquitoes. These results align with the trade-off hypothesis, which states that multi-host viruses that adapt to one host lose fitness in the other hosts [41, 42]. When their neutralization sensitivity was assessed, NS1-T139A escaped neutralization, but E-V355I was more sensitive to neutralization. These results demonstrate that neutralization escape is not necessary for viruses that have been passaged in cross-reactive immune environments. As a whole, these results suggest that pre-existing immunity may play a significant role in ZIKV evolution.

## Methods

### Cells and Viruses

We obtained Vero cells (CCL-81) and U937-DC-SIGN cells (CRL-3253) from the American Type Culture Collection (ATCC). HEK293A cells were kindly provided by Dr. Jamie Smyth from the Fralin Biomedical Research Institute. A549 cells were kindly provided by Dr. Nisha Duggal from the Virginia-Maryland College of Veterinary Medicine. All cells were maintained at 37°C with 5% CO_2_. Vero, HEK293A, and A549 cells were cultured in Dulbecco’s modified Eagle’s medium (DMEM) supplemented with 5% fetal bovine serum (FBS), 1% nonessential amino acids, and 0.1%gentamicin. We cultured U937-DC-SIGN cells in Roswell Park Memorial Institute medium (RPMI-1640) supplemented with 2 mM L-glutamine, 5% FBS, 1% nonessential amino acids, 0.1% gentamicin, and 0.05 mM 2-mercaptoethanol. The ZIKV strain used was derived from an infectious clone of strain PRVABC59 [43] and was rescued and passaged once in Vero cells (p1). DENV1 strain R99142 and DENV2 strain PUO-218 were obtained from the CDC. DENV3 strain BC188/97 (NR-3801) and strain DENV4 703-4 (NR-48801) were obtained from the Biodefense and Emerging Infections Research Resources Repository (BEI Resources). West Nile virus (WNV) Kunjin strain SW28919 and yellow fever virus (YFV) strain 17-D were obtained from the University of Texas Medical Branch World Reference Center for Emerging Viruses and Arboviruses.

### Serum Sources

All human samples were de-identified by their respective sources prior to purchase. We obtained serum samples from convalescent dengue patients from the Dominican Republic who tested positive for DENV and negative for ZIKV by ELISA via Boca Biolistics, LLC. These samples were referred to as anti-DENV patients A-D. The serum from the four anti-DENV patients was subsequently pooled at equal volumes (hereafter called the anti-DENV pool). Convalescent Zika serum was obtained from BEI Resources (NR-50867). Control serum, derived from blood donors from Kentucky, USA, was from Valley Biomedical Products and Services, INC.

### Plaque Reduction Neutralization Test (PRNT)

Serum samples were serially diluted in RPMI-1640 with 10 mM HEPES and 2% FetalPure bovine serum (Genesee Scientific 25-525H), hereafter referred to as viral diluent. Serum samples were mixed with 800 plaque-forming units per mL (PFU/mL) of the virus of interest. The mixture was then incubated at 37°C for one hour, and the virus-serum mixture was used to inoculate wells in a confluent 24-well plate of Vero cells.

After a one-hour adsorption period, plaque assay overlay media was added to each well, as previously described [44]. We fixed the plates four to seven days later, depending on the virus. We defined the PRNT_50_ as the highest reciprocal dilution that neutralized the virus by at least 50%.

### Viral Passaging

Vero cells were plated to an 80-90% confluency in 24-well plates. On the day of the infection, we mixed ZIKV with an equal volume of the appropriate serum. The virus-serum mixtures were incubated for one hour at 37°C and then used to inoculate the Vero cells. After the one-hour adsorption period, the cells were washed in phosphate-buffered saline (PBS); we then added fresh media supplemented with the appropriate human serum. Cells were monitored every 12 hours, and the supernatant was harvested once the cells demonstrated 50-75% cytopathic effect (CPE). Harvested supernatant was stored at −80°C until they were titered plaque assay. This process was repeated for a total of five passages, where the multiplicity of infection (MOI) for each passage was maximized based on the available titer (.01-1). These experiments were performed in triplicate, producing three unique lineages for each serum condition.

### Library Preparation and Next-Generation Sequencing (NGS) Analysis

We prepared libraries of the unpassaged virus, virus passaged in the anti-DENV pool, and virus passaged in the control serum. To enrich for encapsidated virus and to remove nucleic acids, 120 μL of viral supernatant was mixed with 15 μL of 250 units/mL of Benzonase (Millipore Sigma E1014-5KU) diluted in 10x Benzonase Buffer (200 mM Tris-Cl [pH 7.5], 100 mM NaCl, 20 mM MgCl2) [45–47] and 15 μL of 250 units/mL of RNAse A (Millipore Sigma 10109142001) [48]. Samples were then incubated at 37°C for three hours. RNA was extracted from the samples using the Zymo Quick-RNA Viral Kit (R1035). After extraction, we purified the samples using a 0.8x bead selection with the sparQ PureMag magnetic beads (95196-005) to remove small RNA fragments. First-strand cDNA synthesis was performed using random nonamers and the Maxima H-Reverse Transcriptase kit (EP0751). We synthesized second-strand cDNA using the Q5 High-Fidelity 2X Master Mix (M0492S) [49]. NGS Libraries were then produced using the sparQ DNA Frag & Library Prep Kit from Quantabio (Cat. 95194-024). Samples were sequenced using 150 bp paired-end reads on the Illumina Novaseq 6000 (Novogene Co., Ltd.). Bioinformatic analysis was performed as previously described [50, 51]. Briefly, adapters and bases with <Q30 were trimmed using Bbduk [52]; the remaining reads were mapped to the ZIKV reference using the Burrows-Wheeler Aligner (BWA)[53]. Variants were called with LoFreq [54], and new consensus sequences were produced using the Genome Analysis Toolkit (GATK) [55]. To perform diversity and selection analysis, we used SNPGenie [56] to calculate the non-synonymous (π_N_) and synonymous (π_S_) diversity. To determine the divergence of passaged samples, we generated an all-site VCF file using bcftools [57] and calculated D_xy_ using pixy [58].

### Generation and Rescue of Single Mutant Viruses by Bacteria-Free Cloning

Mutagenic PCR primers were designed *in silico* to introduce the identified mutations using SnapGene 6.0.2 software (GSL Biotech). We performed PCRs using the SuperFi II Master Mix (Thermo Fisher 12368010), and the products were gel purified using the Machary-Nagel NucleoSpin Gel and PCR Clean-up kit (740609). We assembled the products using the NEB Builder HiFi DNA Assembly Master Mix (E2621L). The assemblies were digested with DpnI (R0176S), Lambda exonuclease (M0262S), and Exonuclease I (M0293S) and amplified using SuperPhi RCA Premix Kit with Random Primers (Evomics catalog number PM100) [59]. As previously described, RCA products were then transfected into HEK293A cells to produce p0 stocks of the virus [51, 60]. The p0 stocks were then used to infect Vero cells at an MOI of 0.01 to produce p1 stocks, which were used for all downstream tests. The p1 stocks were verified by Sanger sequencing to confirm that the mutant of interest was introduced.

### *In vitro* Competition Assays

Competition assays were performed similarly to previously described methods [51]. Briefly, the mutant and wild-type viruses were mixed at a 1:1 PFU ratio to prepare the competition mixes. This mix was then confirmed by plaque assay, and we amplified a PCR amplicon around the mutation site using extraction-free one-step RT-PCR [61] with qScript XLT One-Step RT-PCR Kit (95143-200). We then purified the amplicon and submitted it for Sanger sequencing, and then assessed the ratio of each virus from the resulting chromatograms using QSVanalyzer [62]. After confirmation, we used the mixes to infect cells at an MOI of 0.01 for each virus. For Vero and A549 cells, virus mixes were added to the cells, and after one-hour adsorption, cells were washed with PBS, and fresh media was added. For the Vero cells with serum supplementation experiments, the mixes were treated identically to the passaging experiments described above. For U937-DC-SIGN cells, we first centrifuged the cells, removed the media, and the cells were resuspended in the diluted virus mix [7]. After a two-hour adsorption period, we washed cells with PBS before adding fresh growth media [7]. Virus was harvested at 2 (U937-DC-SIGN), 3 (Vero and A549), or 4 (Vero with convalescent DENV serum) days post-infection (dpi). Extraction-free RT-PCR and Sanger sequencing were performed on all samples, and the data was analyzed using QSVanalyzer [36]. Relative fitness was calculated as W(t) = F(t)/F(0), where F(t) is defined as the ratio of the mutant virus following the competition and F(0) is defined as the ratio of the mutant virus at baseline [36]. We considered the mutant to have increased fitness in the tested environment if W>1. In contrast, we considered the mutant virus to have reduced or no fitness change if W<1 or W=0, respectively.

### Mosquito Rearing and *In vivo* Competition Assays

We reared *Aedes aegypti* Poza Rica mosquitoes using previously published methods [63]. Briefly, mosquitos were maintained at 28°C with a relative humidity of 75% and a 12:12 (light/dark) photoperiod. During larval stages, mosquitoes were maintained on ground Nishikoi fish food. Adult mosquitoes were given 10% sucrose solution *ad libitum* via cotton balls. Mosquitoes were separated from the colony 6 – 8 days post eclosion at a 5:1 ratio of females to males into disposable 16 oz containers. Mosquitoes were starved of glucose and water for 24 hours before the infectious blood meal of defibrinated sheep’s blood. Mosquitoes were fed for one hour using the Hemotek membrane feeder system (SP4W1-3). After feeding, mosquitoes were anesthetized at 4°C, and females fed to repletion were separated into a new container. We maintained these mosquitoes for ten days under rearing conditions with sucrose *ad libitum*. Mosquitoes were then anesthetized using triethylamine, and whole bodies were collected in viral diluent supplemented with 50 μg/mL gentamicin and 2.5 μg/mL of amphotericin B. Samples were processed by adding a single sterile metallic bead per tube, homogenizing the samples at 30 freq/s for 2 min using the Qiagen TissueLyser II (85300), and diluting the samples 1:5 in viral diluent. We used the extraction-free RT-PCR, Sanger sequencing, and QSVanalyzer workflow described above to analyze the samples. The “naïve” blood meal consisted of defibrinated sheep’s blood, 10^6^ PFU/mL of the competition mix, and 0.5 μM ATP. For the “immune” blood meal, we pretreated the virus for one hour by incubating it with anti-DENV serum, but other conditions were unchanged.

### Statistical Analysis

Statistical analysis was performed in Prism 9 (GraphPad, San Diego, CA, USA). A two-way ANOVA with Šídák’s correction for multiple comparisons was used to compare the divergence and non-synonymous diversity (π_n_) within the antibody binding regions of the passaged populations. A two-way ANOVA with Dunnett’s correction for multiple comparisons was used to compare the neutralization of the mutant viruses to the wild type. For *in vitro* competition assays, a Shapiro-Wilk test for normality was performed to ensure normality, and then a one-sample t-test was performed. For the *in vivo* competition assays, a Fisher’s exact test was used to determine which species (mutant or WT) would become dominant in a population of mosquitoes.

## Results

### Donor Serum Characterization

We first evaluated the serostatus of the donors from the Dominican Republic (referred to as anti-DENV patients A-D) and the US (Kentucky; hereafter referred to as control serum) by performing plaque reduction neutralization tests (PRNTs) against DENV and ZIKV. Data are presented as the reciprocal serum dilution which resulted in a 50% reduction in plaques (PRNT_50_) (Table 1). Since closely related flaviviruses are known to be cross-neutralized by convalescent serum, we defined the infecting virus as having a PRNT50 ≥ 4-fold higher than the other viruses within the group [64, 65]. Based on these parameters, we concluded that all anti-DENV donors had a history of DENV infection without prior ZIKV infection. Furthermore, we concluded that the control serum donors had no history of infection with either DENV or ZIKV.

**Table 1:** Serological Characterization of Donor Blood. Donor blood was serially diluted and mixed with dengue virus (DENV) serotypes 1-4 or Zika virus (ZIKV). The reciprocal of the highest serum dilution that neutralized 50% of the challenge virus is reported as the PRNT50 value. These data represent two biological replicates, each with three technical replicates.

| Virus | anti-DENV<br>Patient A | anti-DENV<br>Patient B | anti-DENV<br>Patient C | anti-DENV<br>Patient D | Control<br>Serum |
| --- | --- | --- | --- | --- | --- |
| DENV1 R99142 | 640 | 1280 | 160 | 10240 | <20 |
| DENV2 PU0218 | 640 | 10240 | 80 | 2560 | <20 |
| DENV3 BC188/97 | 1280 | 320 | 640 | 5120 | <20 |
| DENV4 703-4 | 640 | 640 | 360 | 1280 | <20 |
| ZIKV PRVABC59 | 20 | <20 | <20 | 80 | <20 |

To increase antibody diversity, we pooled the four anti-DENV donors (hereafter referred to as the anti-DENV pool) at equal volumes. We then tested the pool’s neutralization of DENV1-4 and ZIKV (Figure 1), as well as yellow fever virus (YFV) and West Nile virus (WNV) (Supplemental Table 1). The cross-reactive PRNT_50_ of the anti-DENV pool against ZIKV was 40, which represents an intermediate between the specific neutralization from the convalescent ZIKV patient (PRNT_50_ = 320) and the control serum (PRNT_50_ < 20). Thus, we generated an anti-DENV pool with a history of DENV infection but no previous exposure to ZIKV, WNV, or YFV.

**Figure 1:**
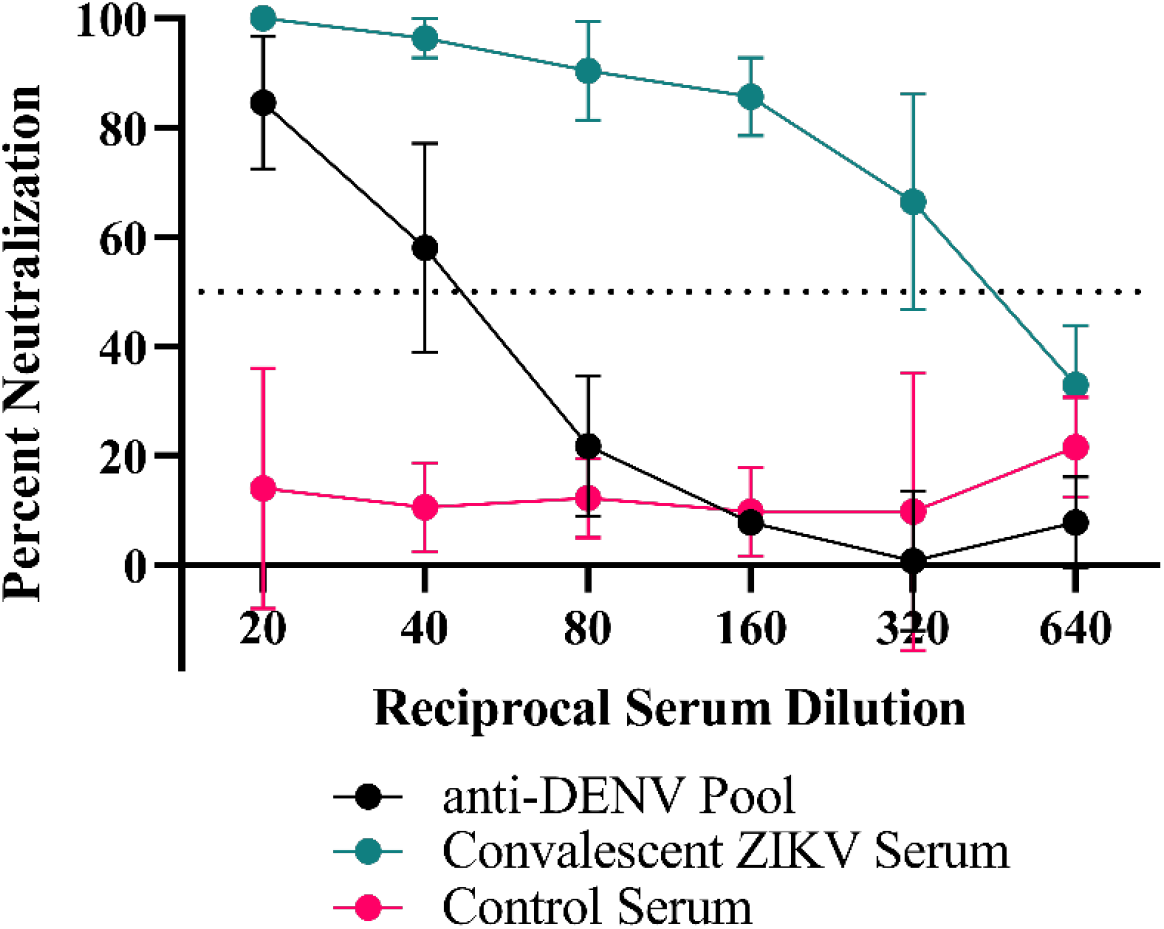
Zika virus (ZIKV) neutralization by the anti-dengue virus (DENV) serum pool. The anti-DENV pool, control serum and convalescent ZIKV serum were serially diluted and mixed with 800 PFU/ml of ZIKV. The reciprocal of the highest serum dilution that neutralized 50% (presented as a dotted line) of the challenging virus is reported as the PRNT_50_ value. These data represent two biological replicates, each with three technical replicates, and the error bars represent the standard deviation from the mean.

### Passaging Virus in the Presence of anti-DENV Serum

To assess the impact of convalescent dengue serum on ZIKV evolution, we adapted previously reported immune-driven evolution protocols [18, 22]. Specifically, we mixed ZIKV at a 1:1 ratio by volume with either the anti-DENV pool at the PRNT_50_ concentration (1:40 dilution) or control serum at the same dilution (Figure 2A). After one hour of incubation, we infected Vero cells with the virus-serum mixtures. After adsorption, we washed the cells and maintained them with media supplemented with anti-DENV pool or control serum. We monitored the cells and harvested the supernatant when we observed >75% cytopathic effect (CPE). We then titered the harvested virus (Figure 2B) and used it for a subsequent passage. By the fifth passage, we observed that the time to produce CPE in the anti-DENV pool passaged virus increased by 1.5 days from the initial passage, while the control serum passaged virus decreased by 0.5 days (Supplemental Figure 1). Given this phenotypic change, we next sequenced the virus populations.

**Figure 2:**
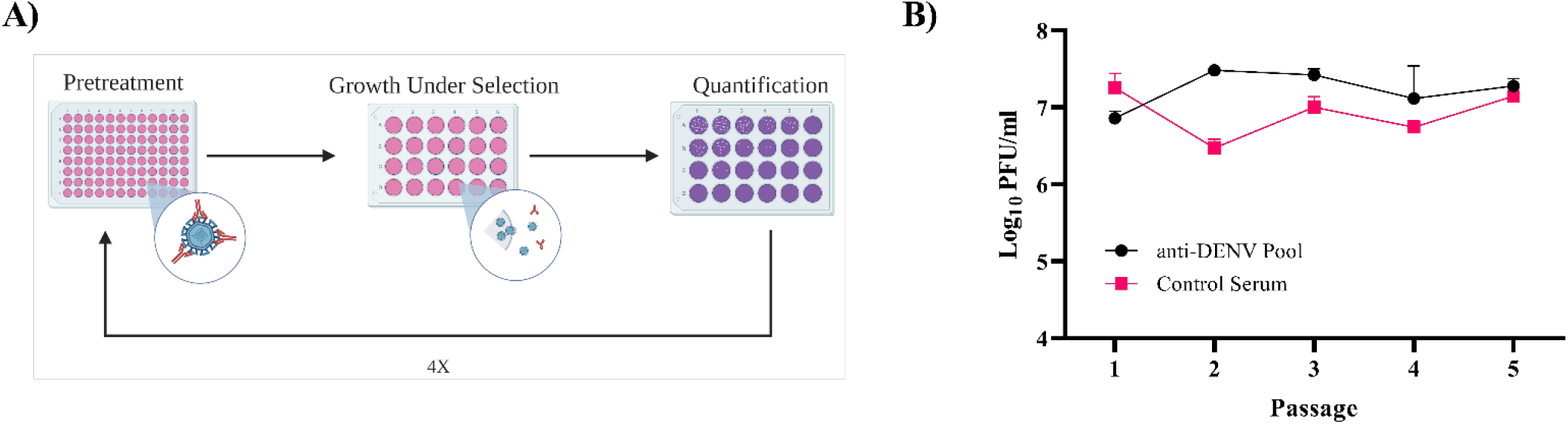
Zika virus (ZIKV) Passaging Workflow. A) Virus was mixed with either anti-dengue virus (DENV) or control serum that had been diluted 40-fold and then incubated for one hour at 37°C. The mix was then used to infect Vero cells. After adsorption and washing, we added fresh media that contained either anti-DENV serum or control serum. Supernatant was harvested when the cells showed at least 75% cytopathic effects. After clarifying and freezing the supernatant, it was assessed by plaque assay to quantify infectious virus. This process was repeated a total of five times. B) The post-passage titers of the anti-DENV pool and control serum passaged virus, as measured by plaque assay.

### Population Divergence, Diversity, and Mutation Selection

We next sought to determine the evolutionary impact of passaging ZIKV in convalescent dengue serum, hypothesizing that the viral populations would differ significantly from the virus passaged in control serum. To this end, we performed Illumina next-generation sequencing (NGS) on viral RNA from the unpassaged virus, the control serum passaged virus, and the anti-DENV pool passaged virus. We included the unpassaged virus to establish an accurate starting consensus sequence. By sequencing the virus passaged in the control serum and the anti-DENV pool, we could differentiate between mutants derived from passaging ZIKV in Vero cells with human sera (control) and the impact of passage in the presence of dengue convalescent serum.

For our analysis, we focused on the regions of known antibody binding: premembrane (prM), envelope (E), and nonstructural protein 1 (NS1) [66]. We examined the average nucleotide distance (divergence) between our unpassaged and passaged viruses (Figure 3A) and the non-synonymous diversity, π_n_ (Figure 3B), within the antibody binding regions. We observed significantly lower divergence (*p* = 0.043) and non-synonymous diversity (*p* = 0.001) in the prM region of the anti-DENV pool passaged virus compared to the control serum passaged virus. These results suggest that selection by anti-DENV serum limited the divergence and diversification of the prM protein.

**Figure 3:**
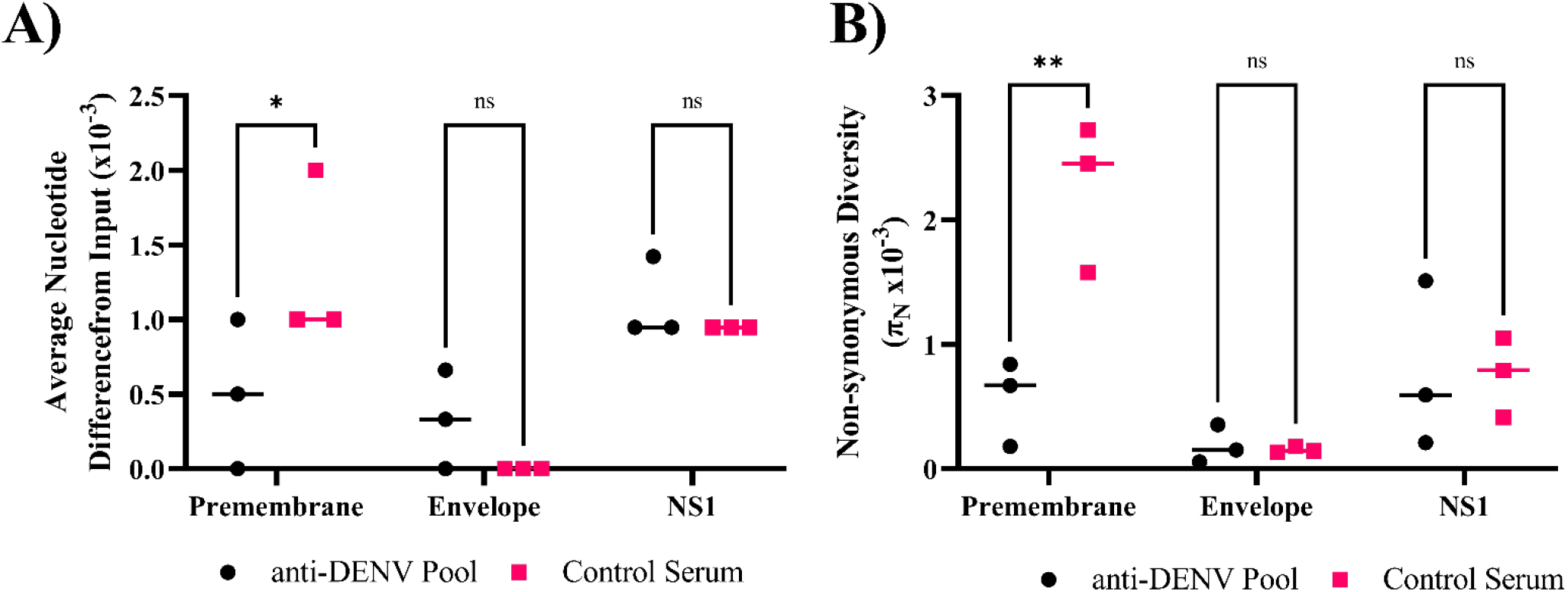
Anti-dengue virus (DENV) Immune Selection Limits Zika virus (ZIKV) Divergence and Nonsynonymous Diversity in the Premembrane Protein. A) Divergence within the antibody binding region of passaged samples compared to the input virus. B) Comparing the πn values at each protein with the antibody binding region of the passaged populations. Statistics were performed using a two-way ANOVA with a Šídák’s correction for multiple comparisons (* p = 0.043 ** p = 0.001). NS – nonstructural protein

We next examined the antibody binding region for single nucleotide variants enriched after passaging in the presence of convalescent dengue serum. To prioritize variants, we used an allelic frequency cutoff of 0.1 as our focus was on higher frequency mutations, which we expected would be more likely to have a phenotypic impact on the virus. Of the 43 mutations identified across the input virus and the passaged populations at an allelic frequency ≥ 0.1 (Supp. Table 2-4), five non-synonymous mutations were found exclusively in the anti-DENV pool passaged virus within the antibody binding region that increased in frequency (Table 2). We selected two mutations to construct within the envelope (E), E-V355I, and NS1, NS1-T139A (Figure 4A), regions which have been shown to mediate cell entry [67] and alter infectivity in mosquitoes [37, 68], respectively. Interestingly, ZIKV E-355I has been previously isolated from a human in Brazil (NCBI Accession OL423668) [69]. We used a bacteria-free cloning approach to generate mutant viruses [70, 71] and a previously characterized ZIKV infectious clone as a template [43].

**Figure 4:**
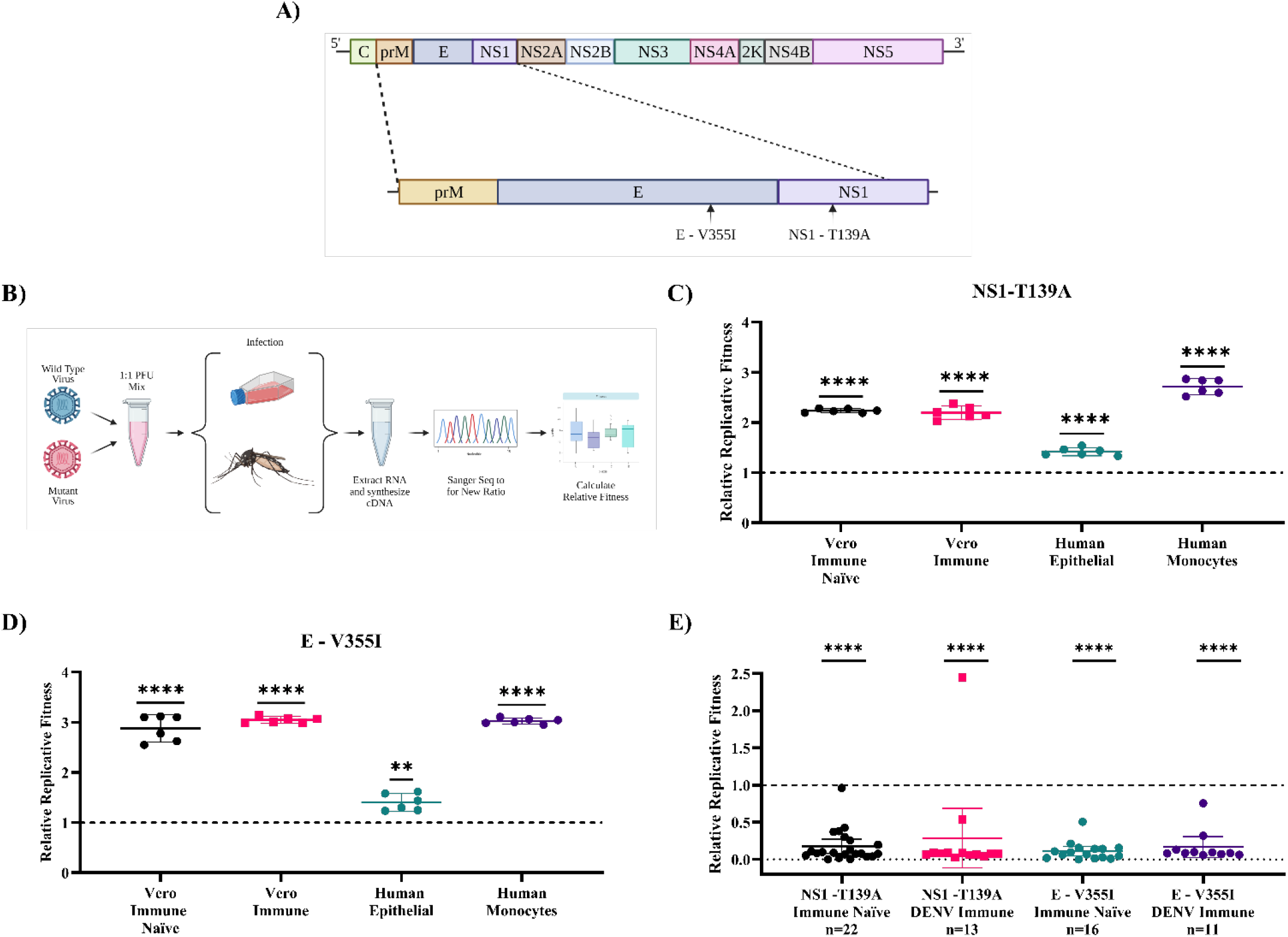
Zika virus (ZIKV) mutants have enhanced fitness in mammalian cells and reduced fitness in mosquitoes. A) Within the ZIKV genome, antibody selection primarily occurs against the M, E, and NS1 proteins. Within those regions, based on several criteria, we selected to generate mutations at E-V355I and NS1-T139A. B) To assess replicative fitness, WT virus was mixed with a given mutant virus at a 1:1 PFU ratio. The mix was then used to infect cell lines (MOI = 0.01) or live mosquitoes (10^6^ pfu/ml). After infection, virus was harvested, RNA extracted, and RT-PCR was performed. The resulting amplicon was sequenced to calculate the relative fitness of each mutant. (C-E) In vitro competition assays. NS1-T139A (C) and E-V355I (D) in Vero cells without supplementation of anti-DENV serum (Immune Naïve), Vero cells with supplementation of anti-DENV serum (Immune), A549 cells (Human Epithelial), and U937-DC-SIGN cells (Human Monocytes) were performed. These data represent two biological replicates and three technical replicates, and the error bars represent standard deviation from the mean. Statistical analysis was performed using a one sample t-test (**** p< .0001 and ** p = .0021) with a null value of 1. E) Competition assays in mosquitoes were performed by feeding Aedes aegypti Poza Rica mosquitoes with either untreated virus (Immune Naïve) or virus pre-treated with anti-DENV serum (DENV Immune). Each data point represents a single mosquito, and the error bars represent the standard deviation from the mean of pooled fitness across all mosquitoes. The pooled group sizes from two independent experiments are reported below the group titles on the x-axis. Statistical analysis was performed using a Fisher’s exact test (**** p < .0001).

**Table 2:** Variants Identified in Passaged Zika virus (ZIKV) Populations. Non-synonymous mutations within the key antibody binding domains of ZIKV (prM, E, and NS1) with an allelic frequency ≥ 0.1. If a mutation was not found in a replicate, the box is grayed out. The mutations that were selected for downstream use are bolded. NS1 refers to nonstructural protein 1.

| nt Position | Protein | Reference | Alternative | AA Change | Allelic Freq. Rep. 1 | Depth Rep. 1 | Allelic Freq. Rep. 2 | Depth Rep. 2 | Allelic Freq. Rep. 3 | Depth Rep. 3 |
| --- | --- | --- | --- | --- | --- | --- | --- | --- | --- | --- |
| 721 | Premembrane | A | G | [H/R] | 0.997832 | 13838 | 0.073241 * | 3823 | 0.148524 | 2168 |
| 1435 | Envelope | T | C | [V/A] |  |  |  |  | 0.736453 | 2030 |
| **2040** | **Envelope** | **G** | **A** | **[V/I]** |  |  | 0.920766 | 3395 |  |  |
| **2904** | **NS1** | **A** | **G** | **[T/A]** | 0.998622 | 13788 |  |  | 0.887506 | 2249 |
| 3222 | NS1 | A | G | [K/E] |  |  |  |  | 0.103163 | 3222 |

### Mutants identified during passage in convalescent dengue serum have increased fitness in mammalian cell culture

To assess the impact of the mutations on replicative fitness, we performed competition assays comparing the mutants to WT ZIKV (Figure 4B). We mixed mutant and wild-type viruses at a 1:1 PFU ratio and then used the mixes to infect several cell lines. To calculate the fitness of the mutant virus in each environment, we compared the relative proportion of the mutant virus pre- and post-infection [36]. If the proportion increased during infection, we concluded the virus had increased fitness; if the proportion remained the same or decreased, we concluded the virus had neutral or decreased fitness, respectively. We tested four cell culture environments: Vero cells without supplementation of anti-DENV serum, representing an immune naïve model (Immune Naïve); Vero cells supplemented with the anti-DENV pool, representing a dengue immune model (Immune); A549 cells representing a human epithelial cell model (Human Epithelial); U937-DC-SIGN cells, representing a human monocyte model (Human Monocyte). These cell lines were selected as they previously were shown to be susceptible to ZIKV infection and represent cell types critical to pathogenesis [7, 72, 73]. The NS1-T139A (Figure 4C) and the E-V355I mutant (Figure 4D) had increased fitness compared to WT ZIKV in all environments tested. These data demonstrate that the ZIKV mutants derived from passaging in the presence of anti-DENV serum have increased fitness in mammalian cells.

### Mutants identified during passage in convalescent dengue serum have reduced fitness in mosquitoes

Because ZIKV is transmitted between mammalian and insect hosts, it is critical to test the fitness of these immune-selected mutants in mosquitoes to assess the likelihood that these mutations would be maintained in nature. To that end, we fed *Aedes aegypti* an artificial blood meal containing the WT/mutant competition mixes used in the previous studies. To mimic the impact of a prior DENV infection, we also performed an experiment where we pretreated the virus with the anti-DENV serum pool before the blood meal. We then collected mosquitoes at ten days post-feeding, and whole-body homogenates were tested to determine the proportion of each virus compared to the original blood meal (Figure 4E). Regardless of host immune status, the mutant viruses had significantly reduced fitness compared to WT ZIKV in mosquitoes (*p*<.0001). These data suggest that the ZIKV mutants derived from cross-reactive immune selection have significantly reduced fitness in the mosquito host. These results, taken with those from the *in vitro* competitions assays, indicate that our findings align with the trade-off hypothesis [41, 42].

### Neutralization escape is not required for fitness enhancement during cross-reactive immune selection

To understand the impact of immune selection on neutralization sensitivity, we performed PRNTs on the ZIKV mutants using the anti-DENV pool (Figure 5). We hypothesized that the mutants would escape neutralization because they were generated from passaging in anti-DENV serum. The anti-DENV pool neutralized wild-type ZIKV at a PRNT_50_ of 40, similar to what we found previously (Figure 1). The mutants showed polarized neutralization susceptibilities; specifically, the anti-DENV pool no longer neutralized NS1-T139A (PRNT_50_ <20), indicative of escape, while E-V355I was slightly more sensitive to neutralization than the WT (PRNT_50_=80 *p* < .0001). Interestingly, when we examined the neutralization sensitivity of the populations from which these mutants were derived, we saw neither escape nor sensitization phenotypes (Supplemental Figure 2). These results suggest that cross-reactive antibody selection can generate mutants with enhanced sensitivity (E-V355I) and reduced sensitivity (NS1-T139A) to antibody selection and that neutralization escape is not necessary for fitness enhancement during cross-reactive immune selection. To further contextualize the results, we generated two mutants from mutations identified in the control population (Supplemental Table 4): prM-S109P and prM-M159V. These mutations were selected because we observed higher genetic divergence and diversity in the prM protein of the control population compared to the anti-DENV pool population (Figure 3A-B). Both mutants from the control serum passaged population showed greatly enhanced sensitivity to neutralization by the anti-DENV pool (PRNT_50_ > 640). This behavior was also observed in the passaged populations that these mutants were derived from (Supplemental Figure 2). These results indicate that passaging in anti-DENV serum protected ZIKV from taking on these extreme neutralization phenotypes. When we assessed the replication of the passaged populations in Vero cells, we observed that the control serum passaged virus had increased replicative fitness (Supplemental Figure 3A). In contrast, when we tested the passaged populations in Vero cells supplemented with the anti-DENV pool, we observed that the control serum passaged virus did not replicate in this environment (Supplemental Figure 3B). These results indicate that cross-reactive immunity constrains the fitness of the population, similar to the constraint in diversificiation and divergence we observed in the NGS data (Figure 2), but also protect the virus from developing extreme neutralization phenotypes.

**Figure 5:**
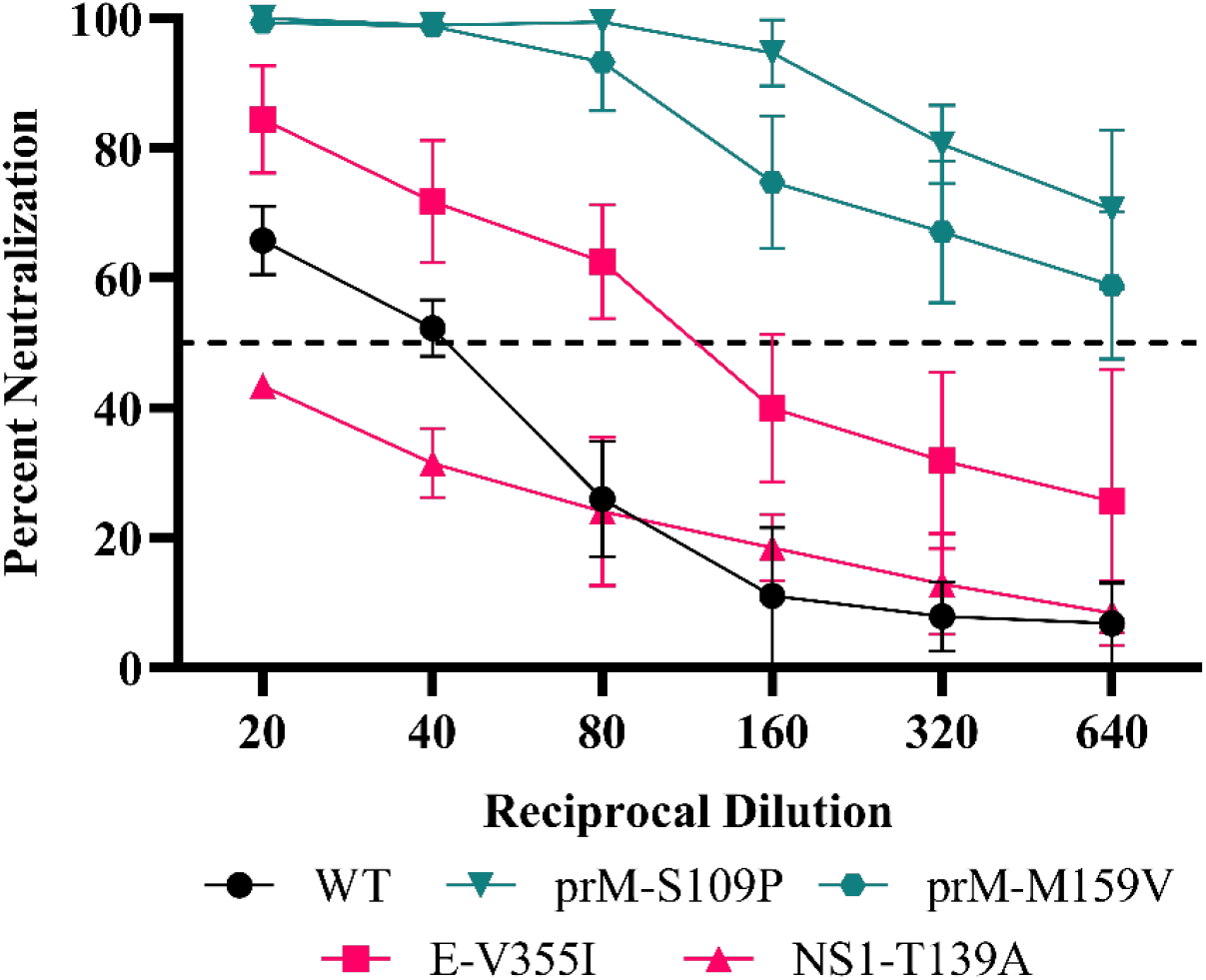
ZIKV Mutant Neutralization Sensitivity. anti-DENV pool were serially diluted and mixed with WT ZIKV (black line), mutants from the anti-DENV pool population (red lines), or mutants identified from the Control Serum-passaged population (green lines). Data are from two biological replicates, each with three technical replicates, and the error bars represent the standard deviation from the mean.

## 1 Discussion

The recent ZIKV outbreak in the Americas was associated with several adaptive mutations that potentially facilitated its rapid emergence [36–40], though the driver of these mutations is unknown. One potential driver of ZIKV evolution is selection mediated by pre-existing adaptive immune responses to closely related flaviviruses such as DENV. While immune-driven evolution has been well-characterized for other viruses [18–25, 28–32], previous studies with ZIKV have some limitations. Regla-Nava et al. sought to examine how pre-existing immunity to DENV affected the evolution and pathology of ZIKV [35]. Using a passaging system in which they oscillated ZIKV infection between mosquito cells and dengue-naïve or dengue-immune mice, they identified a mutant, NS2B I39V, that appeared to escape pre-existing dengue immunity, and increased viral loads, adult and fetal mortality, and mosquito infection [35]. However, NS2B I39V occurred in both the experimental and control populations; therefore, the mutation was likely not driven by immune pressure. Interestingly, we also found NS2B I39V in our anti-DENV passaged population (Supplemental Table 3). Thus, there remained a gap in understanding the specific impact of cross-reactive immune selection on ZIKV evolution. To fill this gap, we passaged ZIKV in the presence of serum from convalescent dengue patients or dengue naïve serum and then studied the evolutionary and fitness consequences on the virus. Within the population passaged in the anti-DENV pool, we identified two mutations, NS1-T139A and E-V355I, that had altered sensitivity to antibody neutralization, increased fitness in mammalian cells, and reduced fitness in *Aedes aegypti* mosquitoes.

Immune selection provides a complex pressure as there is both the enrichment for escape mutations (diversifying selection) within a population resistant to neutralization [17–24, 28–31, 33] and the removal of genotypes sensitive to neutralization (purifying selection) [74]. Two critical aspects of this selection are genetic diversity [75] and divergence [76]: we observed higher genetic divergence and diversity within the prM protein in the control serum-passaged population compared to the virus passaged in the anti-DENV pool. Thus, we concluded that anti-DENV serum likely tempered the ability of ZIKV to diverge during passaging. When we generated two mutants found in the control population, prM-S109P and prM-M159V, we observed that both were highly sensitive to neutralization by the anti-DENV pool, suggesting that the prM protein may be important for ZIKV neutralization by dengue convalescent serum.

Given previous results, we hypothesized that immune-driven evolution would generate neutralization escape mutants [18, 22, 28], as we observed with NS1-T139A; in contrast, E-V355I was more sensitive to neutralization by the anti-DENV pool than WT ZIKV. This possible paradox may be explained by the overall fitness benefit conferred by the E-V355I mutation in all mammalian environments tested here (Fig. 5) [77]. The escape phenotype produced by a nonstructural protein mutant, NS1-T139A, may also appear unexpected since nonstructural proteins are generally not found extracellularly. NS1, however, exists as an intracellular monomer, a membrane-bound dimer, and an *extracellular* hexamer [78]. During infection, secreted NS1 disrupts endothelial junctions, resulting in tissue permeabilization and vascular leakage [79, 80], so neutralizing NS1 may be an important protective strategy. Indeed, passive transfer studies using serum from mice vaccinated with NS1 protected against a lethal ZIKV challenge [81]. These results further demonstrate NS1’s importance as a target for antibody selection. NS1-T139A also increased fitness *in vitro* in mammalian cells, possibly due to its ability to “prime” cells for infection, resulting in increased viral replication [82–84].

While the mutations identified following passage in the presence of anti-DENV serum enhanced fitness in mammalian cell culture, they significantly reduced fitness in live mosquitoes. These results align with the trade-off hypothesis [41, 42], which states that as an arthropod-borne virus, or any virus in a multi-host system, adapts to one of its hosts, it may lose fitness in the other host(s) [41, 42]. These data suggest that while mutants arise following exposure to cross-reactive antibodies in a DENV-immune person, if wild-type virus remains within the population, they might be removed from the population upon mosquito infection, thereby limiting the mutant’s spread.

### Limitations of the study

We used only a single post-epidemic strain of ZIKV and convalescent serum from the Dominican Republic. We may have observed different results using different viruses or serum samples. We focused only on humoral immunity generated from natural infection. However, cross-reactive T cells, which have been shown to play a role in protection against ZIKV [85] and to drive the evolution of other viruses [86–90], should also be examined as a driver of ZIKV evolution. Our passaging approach involved only mammalian cells, representing the multiple rounds of replication within a single host; however, an alternating passaging system between mosquito and mammalian cells could also have been used to mimic the natural transmission cycle [35, 91, 92]. Finally, we generated only single mutants; however, we could have engineered mutant viruses with multiple mutations to assess potential epistatic interactions which would explain the differences observed between the mutant viruses and the populations [93].

### Conclusions

We demonstrated that cross-reactive selection in the presence of anti-DENV antibodies alters ZIKV evolution and fitness. Specifically, we found that passaging ZIKV in mammalian cells with anti-DENV antibodies results in the generation of mutants with altered sensitivity to neutralization. Notably, passaging ZIKV in the presence of anti-DENV serum constrained the evolution of the virus population compared to control populations, which became highly sensitive to neutralization and more fit in Vero cells. However, the mutants from populations passaged in the presence of anti-DENV serum had increased fitness in mammalian cells. This is possibly due to epsistatic interactions reducing fitness. We observed a significant reduction in fitness for both mutants in mosquitoes, consistent with the trade-off hypothesis. These results improve our understanding of the drivers of ZIKV evolution and suggest that future work is needed to more fully dissect the evolutionary implications of inter-flavivirus immune interactions, including inter-serotype interactions with DENV and viruses within the Japanese encephalitis complex (As reviewed [94]).

## Author Contributions

Conceptualization, J.W.L; methodology, J.M.M, and J.W.L; cloning, J.M.M; characterization, J.M.M; formal analysis, J.M.M and J.W.L; validation, J.M.M; writing—original draft preparation, J.M.M; writing—review and editing, J.M.M and J.W.L; visualization, J.M.M, and J.W.L. All authors have read and agreed to the published version of the manuscript.

## Acknowledgments

Figures 2A, 4A, and 4B were generated using BioRender. The Virginia Tech Genomic Sequencing Center performed Sanger sequencing. We thank Christina Chuong and Kelsey Marano for editing the manuscript.

