## Supplemental Data for "Replication in the presence of dengue convalescent serum impacts Zika virus neutralization sensitivity and fitness"

### 1.1 Supplementary Figures

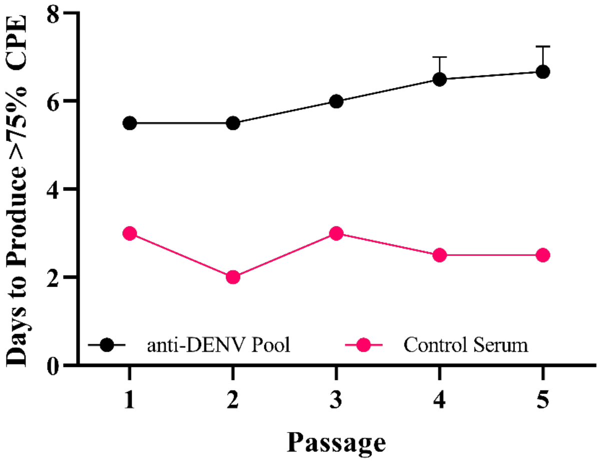


**Supplemental Figure 1:** Cytopathic Effect Production Post Passaging. During passaging, viral supernatant was harvested when >75% of cells demonstrated cytopathic effects (CPE). The days post-infection where this occurred were recorded for each passage. Data represent the three independent lineages within each passaging condition (anti-dengue virus (DENV) Pool or Control Serum).

| Virus | PRNT$_{50}$ Reciprocal Dilution |
|---|---|
| DENV1 R99142 | 10240 |
| DENV2 PUO218 | 2560 |
| DENV3 BC188/97 | 81920 |
| DENV4 703-4 | 5120 |
| West Nile Virus (WNV) Kunjin | 80 |
| Yellow Fever Virus (YFV) 17D | <20 |

**Supplemental Table 1: Serological Characterization of the anti-dengue virus (DENV) Pool.** The anti-DENV pool was serially diluted and mixed with 800 PFU/mL of DENV1-4, West Nile virus (WNV), or yellow fever virus (YFV). The reciprocal of the highest serum dilution that neutralized 50% of the challenge virus is reported as the PRNT50. These data represent two biological replicates, each with three technical replicates.

| nt Position | Protein | Reference | Alternative | AA Change | Allelic Freq. Rep. 1 | Depth Rep. 1 | Allelic Freq. Rep. 2 | Depth Rep. 2 | Allelic Freq. Rep. 3 | Depth Rep. 3 |
|---|---|---|---|---|---|---|---|---|---|---|
| 1436 | Envelope | T | C | [V/V] | 0.089398* | 1745 | 0.102114 | 1939 | 0.114118 | 2550 |
| 2904 | NS1 | A | G | [T/A] | 0.363902 | 2050 | 0.345592 | 1985 | 0.344643 | 2800 |
| 2905 | NS1 | C | T | [T/I] | 0.09196* | 1990 | 0.083739* | 2054 | 0.10833 | 2797 |
| 2922 | NS1 | C | A | [L/I] | 0.40373 | 1984 | 0.415464 | 1940 | 0.402186 | 2745 |
| 10685 | 3’ UTR | G | C | - | 0.148893 | 497 | 0.169043 | 491 | 0.136192 | 793 |
| 10702 | 3’ UTR | A | C | - | | | | | 0.128049 | 164 |

**Supplemental Table 2: Identified Variants in the Unpassaged Zika virus (ZIKV) Populations.** Mutations within the unpassaged ZIKV population with at least one variant with an allelic frequency ≥ 0.1. If a mutation was observed to occur at an allelic frequency < .1, it is marked with a *. If a mutation was not observed at the limit of detection for LoFreq, the box is grayed out. NS refers to nonstructural proteins. UTR refers to the untranslated region.

| nt Position | Protein | Reference | Alternative | AA Change | Allelic Freq. Rep. 1 | Depth Rep. 1 | Allelic Freq. Rep. 2 | Depth Rep. 2 | Allelic Freq. Rep. 3 | Depth Rep. 3 |
|---|---|---|---|---|---|---|---|---|---|---|
| 324 | Capsid | G | A | [G/R] | 0.368546 | 6333 | | | | |
| 721 | Premembrane | A | G | [H/R] | 0.997832 | 13838 | 0.073241 * | 3823 | 0.148524 | 2168 |
| 1435 | Envelope | T | C | [V/A] | | | | | 0.736453 | 2030 |
| 2040 | Envelope | G | A | [V/I] | | | 0.920766 | 3395 | | |
| 2904 | NS1 | A | G | [T/A] | 0.998622 | 13788 | | | 0.887506 | 2249 |
| 3050 | NS1 | T | C | [A/A] | 0.478473 | 14563 | | | | |
| 3222 | NS1 | A | G | [K/E] | | | | | 0.103163 | 3222 |
| 3674 | NS2a | T | C | [A/A] | | | 0.926575 | 3936 | | |
| 4338 | NS2b | A | G | [I/V] | 0.4646 | 13870 | | | | |
| 4502 | NS2b | C | T | [P/P] | 0.454186 | 13915 | | | | |
| 5480 | NS3 | C | T | [F/F] | | | | | 0.712617 | 1712 |
| 5680 | NS3 | C | T | [S/F] | | | | | 0.108287 | 1810 |
| 6336 | NS3 | A | G | [S/G] | | | 0.933468 | 3457 | | |
| 6373 | NS3 | A | T | [K/I] | | | 0.916324 | 4135 | | |
| 6903 | Peptide 2K | T | C | [L/L] | | | 0.919512 | 4510 | | |
| 6954 | NS4b | A | G | [S/G] | 0.998938 | 18839 | | | | |
| 7026 | NS4b | T | C | [S/P] | 0.470057 | 23144 | | | | |
| 7055 | NS4b | A | G | [T/T] | | | | | 0.146478 | 3748 |
| 7463 | NS4b | C | T | [A/A] | | | | | 0.141935 | 1550 |
| 7754 | NS5 | G | A | [K/K] | | | | | 0.31018 | 2505 |
| 7887 | NS5 | T | C | [Y/H] | | | | | 0.737383 | 2239 |
| 8897 | NS5 | A | G | [L/L] | 0.564385 | 11276 | | | | |
| 9227 | NS5 | A | G | [L/L] | | | 0.929367 | 4219 | | |
| 10390 | 3’ UTR | T | C | - | | | | | 0.745545 | 1010 |
| 10680 | 3' UTR | C | T | - | | | 0.268034 | 1414 | | |
| 10685 | 3’ UTR | G | C | - | 0.113841 | 2890 | 0.066581 * | 781 | 0.131336 | 434 |
| 10702 | 3’ UTR | A | C | - | 0.134301 | 551 | 0.116564 | 163 | 0.122449 | 98 |
| 10703 | 3’ UTR | T | C | - | 0.095794 * | 428 | | | 0.113924 | 79 |
| 10766 | 3’ UTR | T | C | - | 0.163636 | 55 | | | | |

**Supplemental Table 3: Identified Variants in the anti-dengue virus (DENV) Pool Passaged Zika virus (ZIKV) Populations.** Mutations within the anti-DENV pool passaged ZIKV with at least one variant with an allelic frequency ≥ 0.1. If a mutation was observed to occur at an allelic frequency < .1, it is marked with a *. If a mutation was not observed at the limit of detection for LoFreq, the box is grayed out. NS refers to nonstructural proteins. UTR refers to the untranslated region.

| nt Position | Protein | Reference | Alternative | AA Change | Allelic Freq. Rep. 1 | Depth Rep. 1 | Allelic Freq. Rep. 2 | Depth Rep. 2 | Allelic Freq. Rep. 3 | Depth Rep. 3 |
|---|---|---|---|---|---|---|---|---|---|---|
| 674 | Premembrane | T | C | [C/C] | 0.545297 | 9813 | 0.708497 | 30082 | 0.856781 | 30003 |
| 798 | Premembrane | T | C | [S/P] | 0.121016 | 7404 | 0.141045 | 23255 | 0.085191 * | 23277 |
| 948 | Premembrane | A | G | [M/V] | 0.64813 | 7085 | 0.787246 | 23177 | 0.868867 | 23724 |
| 950 | Premembrane | G | T | [M/I] | 0.109198 | 7143 | 0.035087 * | 23342 | 0.026538 * | 23815 |
| 2922 | NS1 | C | A | [L/I] | 0.222485 | 8041 | 0.082942 * | 29201 | 0.049856 * | 30107 |
| 3019 | NS1 | T | C | [L/S] | 0.100242 | 9078 | 0.130526 | 32936 | 0.080527 * | 33877 |
| 3674 | NS2a | T | C | [A/A] | 0.546211 | 9013 | 0.716048 | 32759 | 0.85645 | 34448 |
| 6336 | NS3 | A | G | [S/G] | 0.540782 | 7062 | 0.698219 | 27848 | 0.854323 | 27863 |
| 6373 | NS3 | A | T | [K/I] | 0.508521 | 10269 | 0.684632 | 38853 | 0.840522 | 37140 |
| 10702 | 3’ UTR | A | C | - | 0.084746 * | 708 | 0.123345 | 1435 | 0.125207 | 1206 |
| 10703 | 3’ UTR | T | C | - | 0.047934 * | 605 | | | 0.108559 | 958 |
| 10749 | 3’ UTR | T | C | - | 0.119658 | 117 | 0.050157 * | 319 | 0.058036 * | 224 |
| 10766 | 3’ UTR | T | C | - | 0.1 | 60 | 0.154762 | 168 | 0.053435 * | 131 |
| 10771 | 3’ UTR | G | C | - | | | 0.168539 | 89 | 0.191781 | 73 |
| 10773 | 3’ UTR | A | C | - | | | 0.333333 | 39 | | |

**Supplemental Table 4: Identified Variants in the Control Serum Passaged Zika virus (ZIKV) Populations.** Mutations within the control serum passaged ZIKV population with at least one variant with an allelic frequency ≥ .1. If a mutation was observed to occur at an allelic frequency < .1, it is marked with a *. If a mutation was not observed at the limit of detection for LoFreq, the box is grayed out. NS refers to nonstructural proteins. UTR refers to the untranslated region.

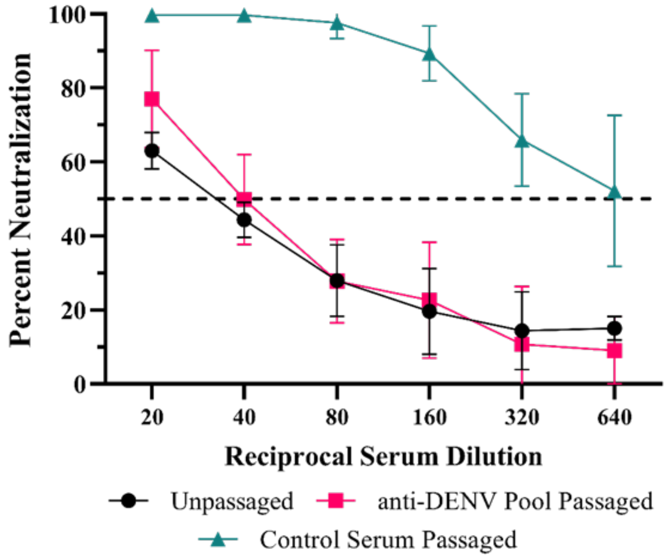


**Supplemental Figure 2: Neutralization Dynamics of Passaged Populations.** The anti-dengue virus (DENV) serum pool was serially diluted and mixed with 800 PFU/mL unpassaged WT ZIKV (black line), passage 5 of the anti-DENV pool population (red lines), or passage 5 of the Control Serum-passaged population (green lines). Data are from two biological replicates, each with three technical replicates, and the error bars represent the standard deviation from the mean.

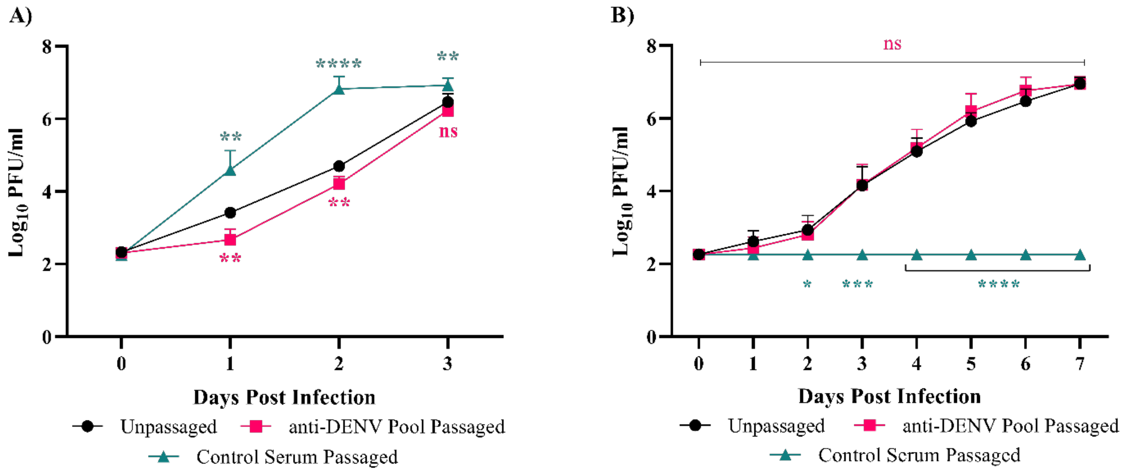


**Supplemental Figure 3: Replicative Fitness of Passaged Populations.** A) Vero cells were infected at an MOI of 0.01 with unpassaged WT Zika virus (ZIKV; black line), passage 5 of the anti-dengue virus (DENV) pool population (red lines), or passage 5 of the Control Serum-passaged population (green lines). B) Vero cells were infected at an MOI of 0.01 with unpassaged WT Zika virus (ZIKV; black line), passage 5 of the anti-dengue virus (DENV) pool population (red lines), or passage 5 of the Control Serum-passaged population (green lines) after the virus was incubated with the anti-DENV pool for 1 hour. Data are from two biological replicates, each with three technical replicates, and the error bars represent the standard deviation from the mean. Statistics were performed using a two-way ANOVA with a Dunnett's correction for multiple comparisons comparing the passaged viruses to the unpassaged virus (ns - nonsignificant, * - p = 0.0139, ** - p = 0.001, *** - p = 0.005, **** - p < 0.0001).
